# Phytoene and phytoene-rich microalgae extracts extend lifespan in *C. elegans* and protect against amyloid-β toxicity in an Alzheimer’s disease model

**DOI:** 10.1101/2024.05.27.595959

**Authors:** Ángeles Morón-Ortiz, Antonis A. Karamalegkos, Paula Mapelli-Brahm, Marina Ezcurra, Antonio J. Meléndez-Martínez

**Affiliations:** Food Colour and Quality Laboratory, Facultad de Farmacia, Universidad de Sevilla, 41012 Sevilla, Spain; School of Biosciences, University of Kent, Canterbury, CT2 7NJ, UK

**Keywords:** Carotenoids, phytoene, bioactive compounds, microalgae, ageing, oxidative stress, antioxidant, anti-ageing, Alzheimer, amyloid-β_42_ proteotoxicity, *Chlorella sorokiniana*, *Dunaliella bardawil*, *C. elegans*

## Abstract

The role of foods is shifting from focusing on the provision of energy and basic nutrients to also include bioactive compounds that prevent the development of chronic age-related diseases. Microalgae offer a source of nutritional compounds with important health effects, and have the potential for production of health-promoting products, without increasing agricultural land use and negatively impacting the environment. Here we investigate the health effects of microalgal extracts with high levels of the colourless carotenoid phytoene. Phytoene is widely available from dietary sources and can be detected across different tissues in the human body. However, it is usually regarded as a precursor for the synthesis of other carotenoids, rather than a bioactive molecule. We utilised the model organism *C. elegans* to show that phytoene-rich extracts from *Chlorella sorokiniana* and *Dunaliella bardawil* have anti-ageing properties. The extracts protect against oxidative damage and amyloid-β_42_ proteotoxicity (a major pathology of Alzheimer’s disease), and extends lifespan. We show that pure phytoene also has these anti-ageing effects, suggesting that phytoene is a bioactive molecule with positive effects on ageing and longevity.

## 1. Introduction

Carotenoids are widespread and versatile compounds that are precursors of derivatives termed apocarotenoids. Both carotenoids and apocarotenoids play key roles in processes that are essential for plant development and resilience (light collection and photoprotection in photosynthesis, communication between plants, pollinators, and seed dispersers, regulation of many plant processes, etc.) and are therefore essential for food security. Carotenoids also function as pigments contributing to the colours of many vegetables and fruits, and as important nutritional components of foods, e.g. as precursors of vitamin A. Many studies indicate that carotenoids have health-promoting properties and contribute to the amelioration or the reduction of the risk of developing diverse diseases (cancer, cardiovascular disease, skin and bone conditions, eye disorders, metabolic disorders, age-related macular degeneration, cognitive impairment, etc.). They can also contribute to skin health, colour and other aesthetic parameters. Therefore, carotenoids are of great interest for the development of a variety of products for human consumption including functional foods, nutraceuticals, supplements, botanicals, cosmeceuticals, or (nutri)cosmetics [1].

The main bulk of dietary intake of carotenoids comes from plant-derived foods (mainly fruits and vegetables), although they are also present in animal-derived foods (e.g., egg yolk, salmon, mussels, dairy), food colorants, and supplements [1]. Microalgae also synthesise carotenoids, as well as many other bioactive compounds with health benefits, and are rich in polysaturated fatty acids and in essential amino acids. Compared to terrestrial plants, cultivation of microalgae is more sustainable due to their rapid growth, ease of cultivation, and non-need for agricultural land, and the commercial importance of microalgae for sustainable production of health-promoting foods is growing. Thus microalgae are a potential source of nutrition that would reduce the need for agricultural land use, decreasing the environmental impact of food production, while improving health [2].

Biosynthesis of carotenoids occurs through the production of phytoene from geranygeranyl pyrophosphate by the enzyme phytoene synthase. Phytoene subsequently undergoes consecutive desaturation and isomerization steps to form lycopene, which is later cyclised to α- and β-carotene. The latter two can be oxygenated to form xanthophylls such as lutein and zeaxanthin. Zeaxanthin is precursor of other typical carotenoids of green microalga, such as violaxanthin and neoxanthin (Figure 1). In contrast to most other carotenoids, phytoene is colourless, does not contribute to the characteristic colours of carotenoid-rich foods and is generally viewed as a precursor without bioactivity in the human body. Phytoene is typically not reported in epidemiological studies and the understanding of its impact on physiology is minimal. However, a few studies have shown that daily intake of phytoene is higher than most other carotenoids, and have detected phytoene in plasma and tissues at concentrations suggesting high levels of bioavailability and bioaccumulation [5–7].

**Figure 1.**
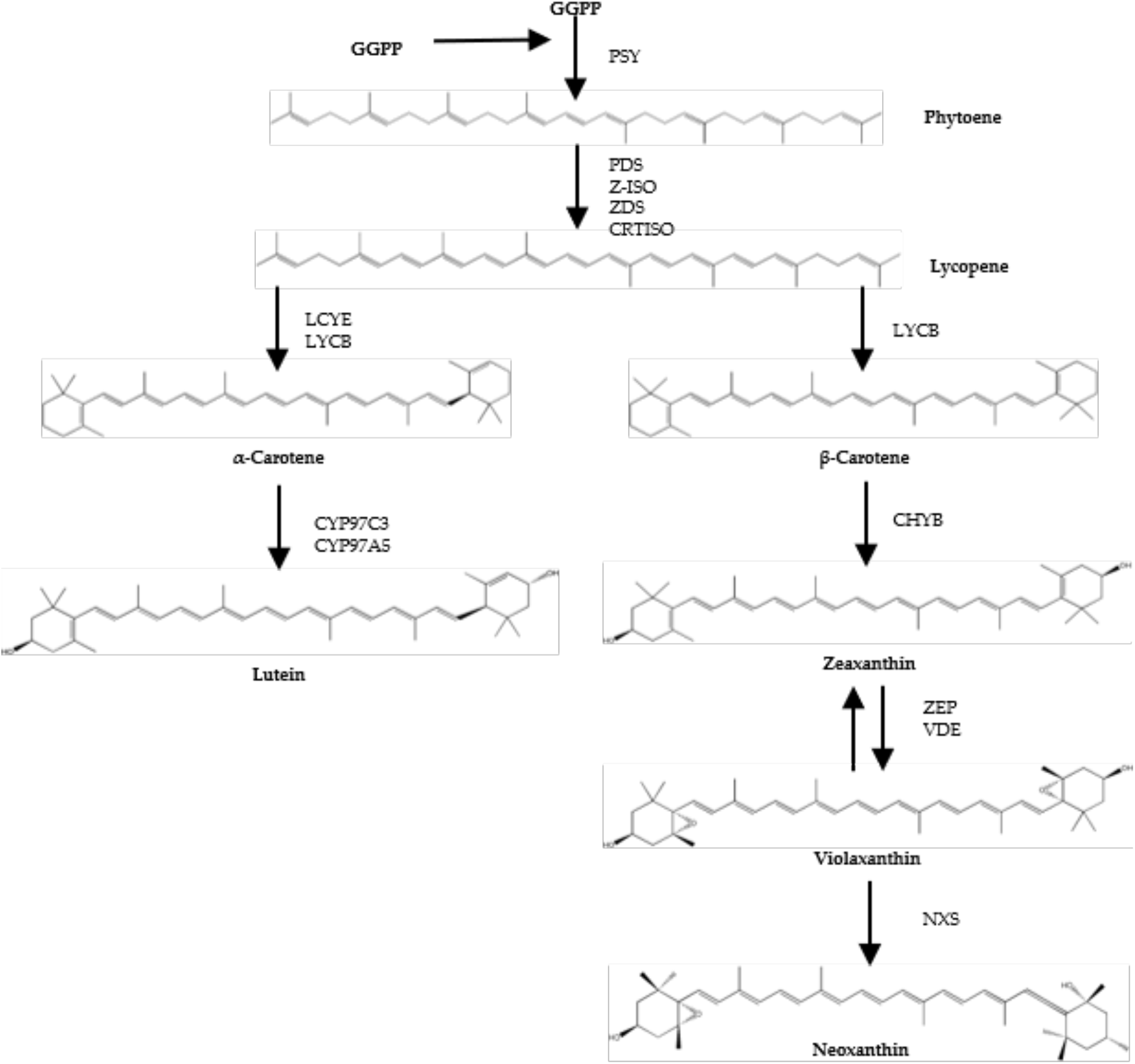
Scheme of biosynthetic steps of microalgal carotenoids (adapted from Valera J.C. et al. (2015) [3], and Lichtenthaler H.K. et al (2012) [4]). GGPP: Geranylgeranyl pyrophosphate; PSY: Phytoene synthase; PDS: Phytoene desaturase; Z-ISO: ζ-Carotene isomerase; ZDS: ζ-Carotene desaturase; CRTISO: Carotene isomerase; LCYE: Lycopene ε-cyclase; LCYB: Lycopene β-cyclase; CYP97C3: Cytochrome P450 ε-hydroxylase; CYP97A5: Cytochrome P450 β-hydroxylase; CHYB: Carotene β-hydroxylase; BKT: β-Carotene oxygenase; ZEP: Zeaxanthin epoxidase; VDE: Violaxanthin de-epoxidase; NXS: Neoxanthin synthase.

Here we examine the bioactivity of phytoene-rich carotenoid extractions from the microalgae *Dunaliella bardawil* and *Chlorella sorokiniana*, and pure phytoene, in the model organism *C. elegans*. We show that both microalgae extracts, as well as phytoene on its own, protect against oxidative stress and amyloid-β toxicity in a model of Alzheimer’s disease, and extend lifespan.

## 2. Materials and Methods

### 2.1 Reagents

Tert-butyl methyl ether (HPLC-grade) and CaCl_2_ were purchased from Honeywell (Seelze, Germany); MeTHF, KH_2_PO_4_, Na_2_HPO_4_, KPO_4_, 5-fluoro-2′-deoxyuridine (FUDR), cholesterol, phytoene standard, and 5-hydroxy-1,4-naphthoquinone (juglone) from Sigma-Aldrich (Steinheim, Germany); methanol (HPLC-grade) and ethyl acetate (HPLC-grade) from VWR Chemicals (Leuven, Belgium); dimethyl sulfoxide (DMSO) was purchased from Duchefa Biochemie (Haarlem, Netherlands); sodium chloride obtained from Fisher Chemical (Hampton, EEUU). Agar, peptone, and LB media from Thermo Fisher Scientific (Hampshire, UK); MgSO4 from Melford (Suffolk, UK).

### 2.2 Microalgae cultivation

#### Chlorella sorokiniana

(211-32) was kindly provided by the algal collection of the Institute of Plant Biochemistry and Photosynthesis (IBVF-CSIC, Seville, Spain) and cultured photo-mixotrophically in liquid Tris-acetate phosphate (TAP) medium [8].

#### Dunaliella bardawil

(UTEX 2538) was supplied from the UTEX Culture Collection of Microalgae (University of Texas, Austin), and cultured using Johnson’s modified medium for *Dunaliella*, as described by Johnson et al. (1968) [9].

### 2.3 Phytoene microalgae enrichment

Cultures of *C. sorokiniana* in the middle of the exponential phase were harvested by centrifugation, resuspended in fresh TAP culture medium, divided into 50 mL-cultures, and incubated with 1 µg/mL of norflurazon, an inhibitor of the carotenoid pathway, to induce accumulation of phytoene [10]. *D. bardawil* cultures were harvested in the middle of the exponential growth phase and incubated with 10 µg/mL of norflurazon [11].

### 2.4 Extraction of carotenoids from microalgae

2-MeTHF, an emerging green solvent that has proved appropriate for the extraction of carotenoids in microalga was used ([10,12]). For extraction, ultrasound-assisted extraction was applied using a frequency of 20 kHz, and amplitude of 30%, and treatment time of 2 min. Samples were centrifuged and the supernatant was transferred to another tube. The procedure was repeated until the sample showed no colour. Subsequently, the samples were concentrated in a rotary evaporator (Eppendorf Concentrator plus™, Eppendorf, Hamburg, Germany), either for use as a worm supplementation or for HPLC analysis.

### 2.5 HPLC analysis

Quantification of carotenoids was carried out using an Agilent (Waldbronn, Germany) 1260 Infinity II Prime LC system. This system was equipped with a diode array detector and a C_30_ column (3 µm, 150 × 4.6 mm) (YMC, Wilmington, NC). Carotenoid extracts were dissolved in 500 µL of ethyl acetate and 10 µL were injected into the system for analysis. Phytoene was detected at 285 nm. The mobile phase consisted of a mixture of methanol, *tert*-butyl methyl ether, and water, delivered at a flow rate of 1 mL/min via a linear gradient, as described in a study by Stinco et al. (2019) [13]. The quantification was performed by external calibration as explained in that study.

### 2.6 Bacterial growth

Bacterial growth curve methodology was implemented in order to ensure no effects of carotenoids in the growth of *Escherichia coli* (OP50). A growth curve was created over 24 hours. The 96-wells microplate assay was employed using OP50 bacteria, and absorbance was measured using a SPECTROstar Nano spectrophotometer (BMG Labtech, Germany). The plate was incubated at 37 °C with shaking for 25 hours. The absorbance was measured at 600 nm every 100 seconds for 900 cycles, and measurements repeated three time for accuracy. Three biological replicates were performed for each condition.

### 2.7 C. elegans culture methods and strains

*C. elegans* were maintained at 20 ° C on Nematode Growth Medium (NGM) plates seeded with *Escherichia coli* OP50. *C. elegans* strains used in this study included: N2 and GMC101 *dvIs100 [unc-54p::A-beta-1-42::unc-54 3’-UTR + mtl-2p::GFP]*, both obtained from Caenorhabditis Genetics Center (CGC, University of Minnesota).

### 2.8 Preparation of experimental plates

35 mm NGM plates were seeded 150 µL OP50 and supplemented with either 50 µL of DMSO control, or 50 µL of 0.2 µg/mL, 1 µg/mL, 2 µg/mL extraction or phytoene standard 48 h later and allowed to dry overnight.

### 2.9 Developmental assays

Thirty adult *C. elegans* were placed on experimental plates and allowed to lay eggs for 2 h. After 52-54 hours, the number of animals at L4 and adult stages was counted to evaluate effects on developmental rate. The experiment was repeated three times.

### 2.10 Oxidative Stress assays

Assays were performed as previously described [14]. Egg lays were performed on experimental plates, approximately 60 h after the egg lay, 50 nematodes at L4 stage were transferred to plates with 400 µM juglone. Survival was recorded hourly over a period of 8 hours. Nematodes were counted as dead if they failed to respond after stimulation with a platinum wire. Any animals that died due to bagging or crawling off the plate were censored. Experiments were carried out in triplicate.

### 2.11 Proteotoxicity assay

The GMC101 strain was used in this assay. Egg lays were performed on experimental plates. Approximately 60 h after the egg lay, animals at L4s stage were transferred to new experimental plates and shifted to 25°C to induce expression of amyloid-β_42_, and kept at 25°C for the remainder of the experiment. Animals were scored every day for four days, and the number of paralysed and non-paralysed animals were counted. Nematodes were counted as paralysed if they were alive but failed to move forwards or backwards after stimulation with a platinum wire. Three biological replicates were performed with three plates containing 30 animals each for each condition.

### 2.12 Lifespan assay

Egg lays were performed on supplemented plates for 2 hours. 60 h later thirty, animals at L4 stage were transferred onto experimental plates treated with 100 µM FUDR. Survival was scored every two days until all animals were dead. Any animals that died due to bagging or crawling off the plate were censored. Three biological replicates were conducted, and each contained four technical replicates.

### 2.13 Statistical analysis

Data processing and statistical evaluation were performed with GraphPad Prism, version 10.1.1 (270) (San Diego, California, USA; www.graphpad.com). Oxidative stress and proteotoxicity assays were analysed using two-way ANOVA. Survival assays were analysed using Kaplan-Meier and Gehan-Breslow-Wilcoxon tests applied for statistical comparison.

## 3. Results

### 3.1 Characterisation of the carotenoid profile of the extracts

To establish if phytoene has effects on health and ageing, we first prepared phytoene-rich extractions from the microalgae *C. sorokiniana* and *D. bardawil*. Phytoene enrichment was induced in *C. sorokiniana* and *D. bardawil* cultures by the addition of norflurazon. We extracted carotenoids from the cultures using the green solvent 2-methyltetrahydrofuran and ultrasound-assisted extraction, as this provides a sustainable extraction method while still giving a high yield. Carotenoid content was quantified by HPLC analysis and showed high levels of phytoene present in the extracts. Phytoene constituted 47% and 45% of the total carotenoid content in *C. sorokiniana* and *D. bardawil* extracts, respectively (Table 1). The *C. sorokiniana* and *D. bardawil* extracts, and a phytoene standard, were used to test for bioactivity throughout this study.

**Table 1.** Concentration (µg/g DW) of carotenoids in *C. sorokiniana* and *D. bardawil* extracts.

| Carotenoid | *C. sorokiniana* | *D. bardawil* |
| --- | --- | --- |
| Lutein | 119.44±5.63 | 77.94±8.52 |
| Zeinoxanthin | 1.79±0.18 | 2.08±0.39 |
| Antheraxanthin | 4.76±0.46 | 3.17±0.29 |
| Zeaxanthin | n.d. | 14.38±1.52 |
| α-Carotene | 6.61±0.6 | n.d. |
| β-Carotene | 12.12±1.14 | 12.78±0.42 |
| (9Z)-β-Carotene | 1.77±0.12 | 1.66±0.22 |
| (15Z)-Phytoene | 117.86±10.44 | 99.68±2.4 |
| (All-E)-Phytoene | 5.05±0.43 | 12.01±0.65 |
| TCC | 259.91±25.37 | 246.34±21.14 |

*n.d.*: not detected. TCC: Total Carotenoid Content.

### 3.2 Effects of phytoene-rich extracts on growth of E. coli OP50

Before performing tests in *C. elegans*, we wanted to rule out the possibility that the extracts inhibit growth of the *C. elegans* food source *E. coli* OP50, as this could lead to reduced food availability and dietary restriction. We conducted bacterial growth assays, treating OP50 bacteria with *C. sorokiniana* and *D. bardawil* phytoene-rich extracts, and with phytoene, at concentrations of 0.2 µg/mL, 1 µg/mL and 2 µg/mL. OD_600_ of the bacterial cultures were measured and used to generate growth curves. OP50 growth was not affected by any of the concentrations (Figure 2), showing that treatment with the phytoene-rich extracts, or with phytoene, does not affect the amount of food available to *C. elegans*.

**Figure 2.**
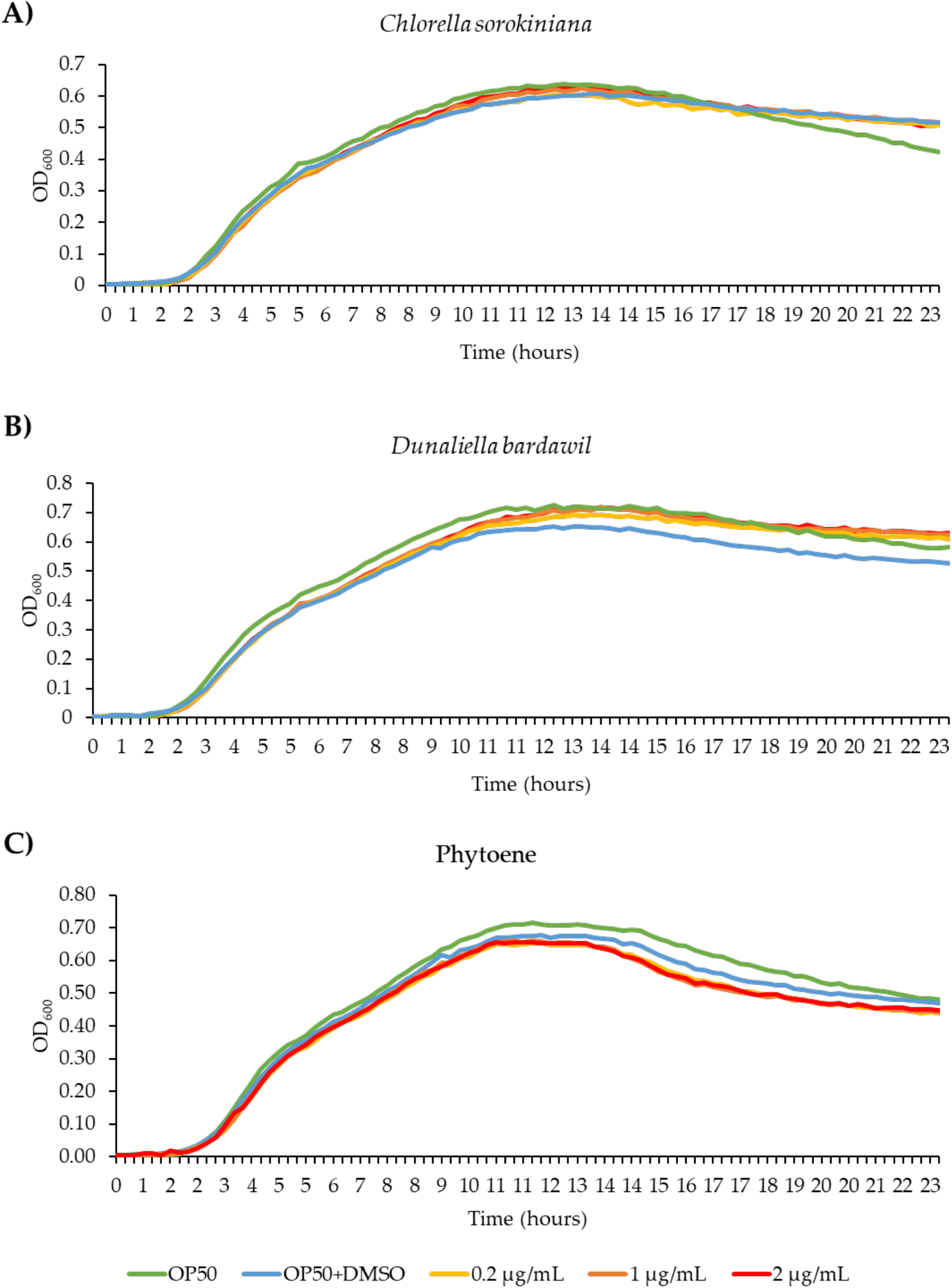
Phytoene-rich extracts and phytoene do not alter growth of the *C. elegans* food source *E. coli* OP50. a-c) Bacterial growth curves (OD_600_) of *E. coli* OP50 cultivated with 0.2 µg/mL, 1 µg/mL, and 2 µg/mL *C. sorokiniana* extracts (a), *D. bardawil* extracts (b) and phytoene standard (c). Three biological replicates were performed.

### 3.3 Effects of phytoene-rich extracts on development of C. elegans

Because natural compounds can have toxic effects, we next asked if the phytoene-rich extracts have detrimental effects on *C. elegans* growth and development. We conducted developmental assays to detect any effects on the development of larvae into adulthood. Extracts at concentrations of 0.2 µg/mL, 1 µg/mL, and 2 µg/mL were tested and no differences were found between the extracts and the controls. Phytoene at the same concentrations did not affect development either (Figure 1), showing that phytoene-rich extracts and phytoene do not exhibit any major toxic effects at the tested concentrations.

### 3.4 Phytoene and phytoene-rich extracts protect against oxidative stress

Next, we asked if the extracts have beneficial effects on health of the animal. Carotenoids are known to have antioxidant properties due to mechanisms including quenching, scavenging, or modulation of gene expression [1]. We therefore tested if the phytoene-rich extracts affect sensitivity to oxidative stress in *C. elegans* using juglone, a naturally occurring mitochondrial toxin that generates superoxide anion radicals [15,16]. We exposed animals to 400 µM juglone and compared survival after 8 hours. We found that in comparison to control conditions, supplementation with the extracts at 0.2 µg/mL) did not affect survival. In contrast, at supplementation with concentrations of 1 and 2 µg/mL, the survival rate of *C. elegans* was increased by 39-53%, depending on the extract and the concentration (*p* < 0.05) (Figure 3). Phytoene only also increased survival at the same concentrations, and to a similar extent, as the *C. sorokiniana* and *D. bardawil* extracts. These findings show that phytoene-rich microalgae extracts, as well as phytoene on its own, have antioxidant properties leading to increased resistance to oxidative stress *in vivo*.

**Figure 3.**
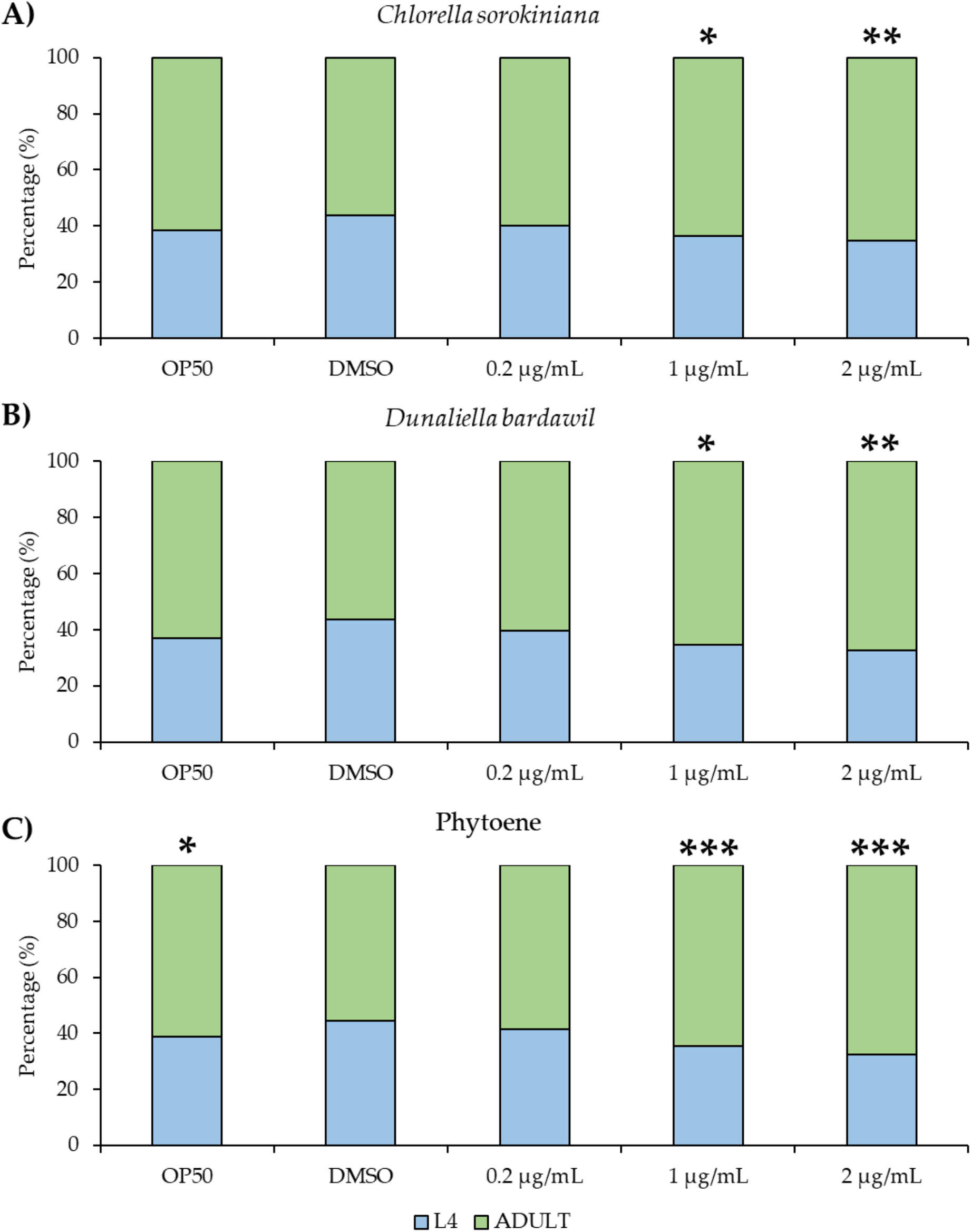
Phytoene-rich extracts and phytoene do not alter development of *C. elegans* (a-c) Percentage of animals that have reached L4 (larval, pre-adulthood) stage and adulthood. Animals were treated with 0.2 µg/mL, 1 µg/mL and 2 µg/mL *C. sorokiniana* extracts (a), *D. bardawil* extracts (b) and phytoene standard (c). Three biological replicates, n=200-300 per condition. ^*^, *p* < 0.05; ^**^, *p* < 0.01; ^***^, *p* < 0.001; ns: Not significant with two-way ANOVA compared to DMSO

**Figure 3.**
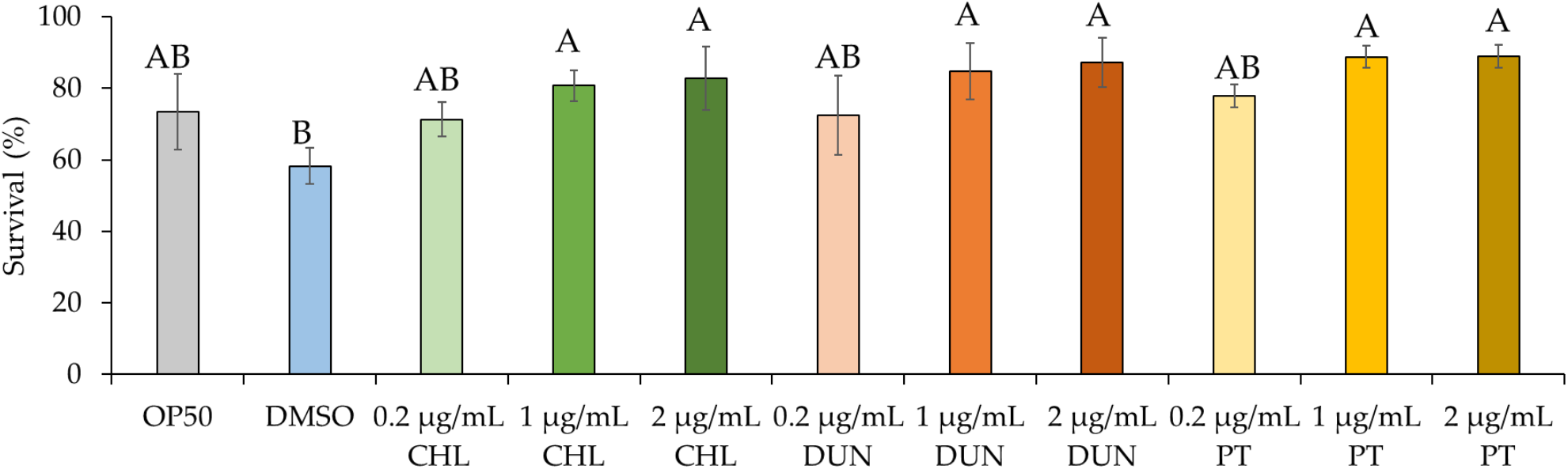
Phytoene-rich extracts and phytoene increase resistance to oxidative stress. Survival after exposure to 400 µM juglone scored after exposure for 8 hours shown. CHL: *C. sorokiniana*; DUN: *D*. bardawil; PT: Phytoene. Data are represented as mean ± SD from three independent experiments with total number of animals n>150 for each condition. Capital letters indicate statistically significant differences (*p* < 0.05) between conditions with two-way ANOVA.

### 3.5 Phytoene and phytoene-rich extracts protect against proteotoxicity

To further investigate protective effects of microalgae extracts on health, we used a humanised *C. elegans* model of proteotoxicity expressing human amyloid-β_42_. In this model, amyloid-β_42_ leads to protein aggregation and formation of amyloid plaques, the main pathology in Alzheimer’s disease. Because the amyloid-β_42_ is expressed in body-wall muscle, protein aggregation leads to muscle dysfunction and paralysis of the animal. We evaluated proteotoxicity by measuring paralysis in animals supplemented with microalgae extracts at 1 µg/mL. Compared to control, supplementation with the extracts, and with phytoene, had a protective effect. On day 2 of the experiment, paralysis was reduced from 62% in the DMSO control to 37%, 40%, and 43% for treatments with *D. bardawil* and *C. sorokiniana* phytoene rich-extracts, and phytoene standard, respectively (*p* > 0.05) (Figure 4). Thus, phytoene protects *C. elegans* against proteotoxicity resulting from human amyloid-β_42_.

**Figure 4.**
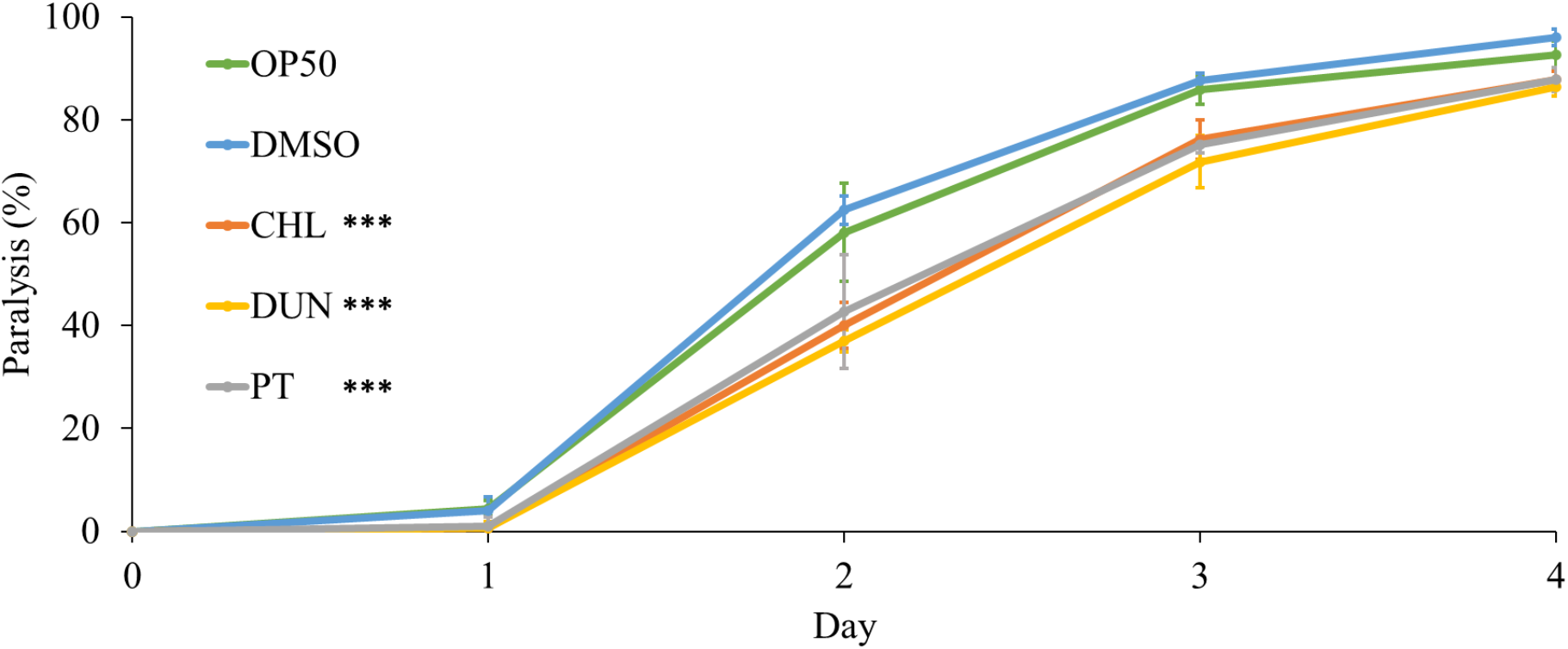
Phytoene and phytoene-rich extracts increase resistance to proteotoxicity. Paralysis of *C. elegans* expressing amyloid-β_42_. CHL: *C. sorokiniana* [1 µg/mL]; DUN: *D. bardawil* [1 µg/mL]; PT: Phytoene [1 µg/mL]. Data are represented as mean ± SD from three independent experiments with total number of animals n>90 for each condition. ^*^, *p* < 0.05; ^**^, *p* < 0.01; ^***^, *p* < 0.001; ns: Not significant with two-way ANOVA.

### 3.6 Phytoene and phytoene-rich extracts extend lifespan

Both oxidative stress and proteotoxicity play central roles in ageing, and the protective effects we observed suggest that phytoene might affect ageing and longevity. We tested this by supplementing animals with the microalgae extracts and with phytoene at 1 µg/ml and conducted lifespan assays. Animals on OP50 had a median lifespan of 16.4 days, similarly to animals on DMSO, which had a median lifespan of 16.1 days. Animals supplemented with extracts from *C. sorokiniana* and *D. bardawil* had median lifespans of 17.7 and 19.1 days, an increase of 10% and 18.6%, respectively. Phytoene also extended lifespan, animals treated with phytoene had a median lifespan of 18.6 days and increase of 15.5% (Figure 5 and Table S1). Our findings suggest that supplementation with phytoene, and phytoene-rich extracts from microalgae, results in lifespan extension in *C. elegans*.

**Figure 5.**
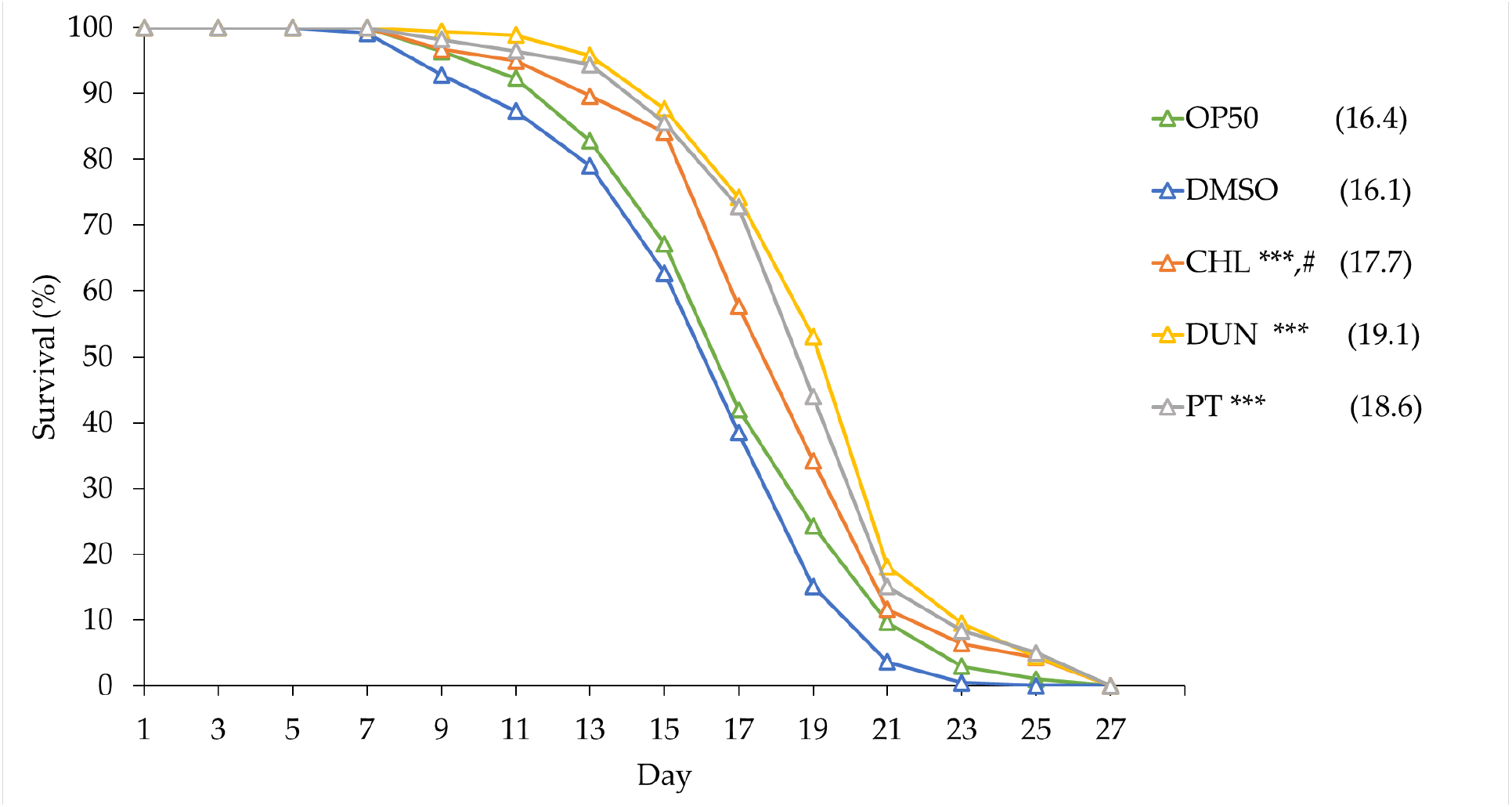
Phytoene-rich extracts and phytoene extend lifespan. Survival curves of *C. elegans* treated with phytoene and phytoene-rich extracts. CHL: *C. sorokiniana* [1 µg/mL]; DUN: *D. bardawil* [1 µg/mL]; PT: Phytoene [1 µg/mL]. Data represented as composites of three replicates, with median lifespan indicated in parenthesis. Total number of animals n>120 for each condition. ^*^, *p* < 0.05; ^**^, *p* < 0.01; ^***^, *p* < 0.001; ns: Not significant with Kaplan-Meier and Gehan-Breslow-Wilcoxon tests.

## 4. Discussion

### 4.1 Insights from analysing bioactivity of phytoene

In this study we have described how phytoene, and phytoene-rich extracts from microalgae, affect resistance to oxidative stress, resistance to amyloid-β_42_ proteotoxicity, and lifespan in *C. elegans*. Our findings show that microalgae are a source of health-promoting compounds and have potential to be developed into functional foods and other products promoting health. Our study also suggests that phytoene is not only a precursor for the biosynthesis of other carotenoids, but also has biological activity in its own right, with beneficial effects on health and longevity. In our study we showed that treatment with phytoene results in anti-ageing effects to the same extent as extracts containing a mixed pool of carotenoids. To our knowledge, no other studies have directly shown that phytoene intake promotes longevity.

Epidemiological studies show that greater intake of foods rich in carotenoids is linked to improved health outcomes. For example, tomatoes are associated with reduced risks of cardiovascular disease and cancer, with these health effects mostly attributed to lycopene. Tomatoes also have high levels of phytoene, but measures of phytoene are not included in most epidemiological studies [17,18], despite some studies showing that phytoene intake is higher than other carotenoids traditionally studied in relation to health, such as lutein, zeaxanthin or β-cryptoxanthin [5,6]. Including a wider range of measures in future epidemiological studies would uncover links between phytoene and health outcomes, and determine if lycopene is the main bioactive component in foods such as tomatoes, or a marker for consumption of foods containing other health-promoting phytochemicals.

### 4.2 Mechanisms by which phytoene impacts ageing

By which mechanisms does phytoene protect against oxidative stress, proteotoxicity and ageing? Carotenoids have antioxidant effects that have been well described in plants, as well as in a large number of *in vitro* and *in vivo* experiments with human relevance [19]. Carotenoids protect against oxidative damage caused by reactive free radicals, including in disease states where oxidative damage occurs such as cancer, diabetes, atherosclerosis, obesity, arthritis and neurodegeneration [20][21]. As already mentioned, carotenoids can act as antioxidants by directly quenching or scavenging reactive oxygen species or through the induction of the expression of genes [1]. In a study the free radical scavenging properties of lycopene, phytofluene, and phytoene were evaluated experimentally and in silico. It was concluded that, although lycopene was the best antiradical, phytoene and phytofluene exhibited a higher antioxidant capacity than could be expected from their fewer number of conjugated double bonds (3 and 5, respectively, compared to 11 for lycopene) [22].

There is growing evidence supporting that carotenoids in addition to directly interacting with reactive oxygen species, also interact with cellular targets, leading to broader effects mediated by endogenous proteins [19]. For example, lycopene and astaxanthin have been shown to inhibit the NF-κB pathway, reducing the activation of downstream pro-inflammatory genes and inflammation caused by obesity, cancer, and other diseases [23]. β-carotene, lycopene, and astaxanthin activate the transcription factor NRF2, resulting in increased expression of antioxidant and detoxification enzymes. Activation of these endogenous defence systems leads to a reduction of intracellular reactive oxygen species, carcinogens and other toxins, and may reduce progression of cancers, cardiovascular diseases, and neurodegeneration [19]. Future studies will show if phytoene also has bioactive properties beyond scavenging reactive oxygen species.

*In vivo* studies in model organisms have directly demonstrated that carotenoids protect against ageing. Several studies used *C. elegans* and showed that supplementation with carotenoid-rich extracts from plant sources not only protect against oxidative stress but also increase lifespan [20,24–29]. Moreover, individual carotenoids improve health in *C. elegans*. e.g. astaxanthin protects against oxidative stress [30] and extends lifespan [31–33] and lycopene protects against amyloid-β_42_ proteotoxicity [26]. Several studies in *C. elegans* also confirmed that the anti-ageing effects of carotenoids require cellular targets and do not only act by directly quenching free radicals. Protection against oxidative stress by carotenoid-rich supplementation is dependent of the Nrf2 orthologue SKN-1 [28]. Lifespan extension by supplementation with Vitamin A is also dependent on the Nrf2/SKN-1 pathway [34], suggesting an important role for Nrf-2/SKN-1 in lifespan extension by carotenoid supplementation. However, individual carotenoids may act through different cellular pathways. Interestingly, astaxanthin promotes longevity in *C.elegans* not through Nrf2/SKN-1, but requires the transcription factor DAF-16, homolog of mammalian FoxO, a critical longevity factor across species [35]. One possibility is that phytoene also stimulates longevity pathways by activation of cellular targets that either overlap, or do not overlap, with those of other carotenoids. This warrants further investigation.

### 4.3 Microalgae – a sustainable source of bioactive compounds to improve healthy ageing?

The role of foods is shifting from focusing on the provision of energy and basic nutrients to also include the supply of bioactive compounds capable of offering protection against the development of diseases. With the global population increasing, and demographics rapidly shifting towards ageing societies with increasing numbers of people suffering from multiple age-related chronic diseases, there is a growing demand for production of foods that meet our growing health needs, while being sustainable [36]. Microalgae are a heterogeneous group of photosynthetic microorganisms with enormous ecological importance that can be harnessed for the sustainable production of foods or ingredients. Thus, they account for approximately half of the global CO_2_ fixation, their cultivation does not require arable land and they grow 10-50-fold faster than plants. Moreover, they are adapted to diverse environments and can thrive in abiotic stressing conditions, including high and low temperatures, nutrient depletion, high salinity or the presence of pollutants. In addition, microalgae have lower nutritional and water requirements compared to terrestrial plants, no requirements of herbicides or pesticides, cultivation produces less wastewater and modulation of biosynthesis of diverse metabolites is easier. Their nutritional value (high quality proteins, unsaturated fatty acids, minerals, vitamins, bioactives) is remarkable, being used as nutritional supplements in aquaculture, human foods, or for the obtaining of food, pharmaceutical, or cosmetic ingredients [2]. In our study we used phytoene-rich cultures of the microalgae species *Chlorella sorokiniana* and *Dunaliella bardawil* and sustainable extraction methods to demonstrate the potential of microalgae as sources for functional ingredients that promote healthy ageing, while reducing land use and increasing sustainability of food production.

## 5. Conclusions

Our study aimed to evaluate the effects of phytoene, and phytoene-rich microalgae extracts, on health in the model organism *C. elegans*. Supplementation with 0.2, 1 and 2 µg/mL of phytoene and phytoene-rich extracts did not affect development, suggesting absence of toxic effects and dietary restriction. Supplementation with 1 and 2 µg/mL, but not 0.2 µg/mL, of the extracts and pure phytoene increased resistance to juglone, a mitochondrial toxin that generates superoxide anion radicals, by 39-53%, showing that phytoene, like other carotenoids, protects against oxidative stress. To assess the effects of phytoene in the context of age-related health, we used a humanised *C. elegans* model of amyloid-β_42_ toxicity, in which in amyloid plaques, a landmark of Alzheimer disease, are formed. We found that 1 µg/mL phytoene and phytoene-rich extracts reduced proteotoxic effects by 30-40%. Finally, we tested the effects of phytoene on lifespan and found that 1 µg/mL phytoene and phytoene-rich extracts increased lifespan by 10-18.6%. Together our results demonstrate that phytoene is a bioactive compound with anti-ageing effects that promote longevity.

## Supplementary Materials

The following supporting information can be downloaded at: www.mdpi.com/xxx/s1, Table S1: Lifespan data of *C. elegans* treated with *C. sorokiniana* and *D. bardawil* extracts and phytoene.

## Author Contributions

Conceptualization, A.J.M.M. and M.E.; methodology, M.E.; investigation, A.M.O., A.A.K. and P.M.B.; resources, A.J.M.M. and M.E; data curation, A.M.O.; writing—original draft preparation, A.M.O.; writing—review and editing, M.E. and A.J.M.M.; supervision, P.M.B.; project administration, A.J.M.M. and M.E.; funding acquisition, A.J.M.M. and M.E. All authors have read and agreed to the published version of the manuscript.

## Acknowledgments

The authors are very grateful to Drs. Rosa León-Bañares and Antonio León-Vaz for the obtaining and provision of phytoene-rich microalgal biomass.

