## Supplemental Table 1 for "Phytoene and phytoene-rich microalgae extracts extend lifespan in *C. elegans* and protect against amyloid-β toxicity in an Alzheimer’s disease model"

| Compound/ Extract | Temp. | FUDR (µM) | Deaths/ censored | Mean lifespan (days) | Lifespan change (%) | *p* vs control |
| --- | --- | --- | --- | --- | --- | --- |
| Control | 20˚C | 100 | **[1-4] 279/81** | **16.05** |  |  |
|  |  |  | [1] 53/7 | 16.36 |  |  |
|  |  |  | [2] 75/15 | 17.24 |  |  |
|  |  |  | [3] 77/13 | 16.15 |  |  |
|  |  |  | [4] 74/46 | 14.48 |  |  |
| *C. sorokiniana* | 20˚C | 100 | **[1-4] 111/130** | **17.66** | **+10.02** | **<0.0001** |
|  |  |  | [1] 34/26 | 20.10 | +22.82 | 0.0014 |
|  |  |  | [2] 56/34 | 17.97 | +4.23 | <0.0001 |
|  |  |  | [3] 10/21 | 17.25 | +6.80 | <0.0001 |
|  |  |  | [4] 11/49 | 16.73 | +15.58 | <0.0001 |
| *D. bardawil* | 20˚C | 100 | **[1-4] 110/189** | **19.18** | **+19.50** | **<0.0001** |
|  |  |  | [1] 32/28 | 20.79 | +27.05 | 0.0015 |
|  |  |  | [2] 23/36 | 19.52 | +13.26 | <0.0001 |
|  |  |  | [3] 30/60 | 18.31 | +13.34 | <0.0001 |
|  |  |  | [4] 25/65 | 19.07 | +31.76 | <0.0001 |
| Phytoene | 20˚C | 100 | **[1-4] 91/123** | **18.58** | **+15.79** | **<0.0001** |
|  |  |  | [1] 28/35 | 21.30 | +30.17 | 0.0026 |
|  |  |  | [2] 32/28 | 18.60 | +7.88 | <0.0001 |
|  |  |  | [3] 23/37 | 17.93 | +10.98 | <0.0001 |
|  |  |  | [4] 8/23 | 18.00 | +24.34 | <0.0001 |

**Table S1. Lifespan data of C. elegans treated with *C. sorokiniana* and *D. bardawil* extracts and phytoene**. 1-4 shows combined data from all trials.
